# Inhibitory engrams in perception and memory

**DOI:** 10.1101/117085

**Authors:** Helen Barron, Tim P. Vogels, Timothy Behrens, Mani Ramaswami

## Abstract

Nervous systems use excitatory cell assemblies or “perceptual engrams” to encode and represent sensory percepts. Similarly, synaptically connected cell assemblies or “memory engrams” are thought to represent memories of past experience. Multiple lines of recent evidence indicate that brain systems also create inhibitory replicas of excitatory engrams with important cognitive functions. Such matched inhibitory engrams may form through homeostatic potentiation of inhibition onto postsynaptic cells that show increased levels of excitation. Inhibitory engrams can reduce behavioral responses to familiar stimuli thereby resulting in behavioral habituation. In addition, by preventing inappropriate activation of excitatory memory engrams, inhibitory engrams can make memories quiescent, stored in a latent form that is available for context-relevant activation. In neural networks with balanced excitatory and inhibitory engrams, the release of innate responses and recall of associative memories can occur through focussed disinhibition. Understanding mechanisms that regulate the formation and expression of inhibitory engrams *in vivo* may help not only to explain key features of cognition, but also to provide insight into transdiagnostic traits associated with psychiatric conditions such as autism, schizophrenia and post-traumatic stress disorder (PTSD).

---

Percepts and memories are thought to be represented in the brain by excitatory activity in groups of neurons, described as cell-assemblies (Hebb, 1949) or memory engrams (Semon, 1921). Excitatory activity within a cell-assembly thus allows percepts to be represented, while coordinated activity across different cell-assemblies faciliates formation and recall of associative memories. These excitatory engrams are stored in quiescent form that allows them to be reactivated in appropriate contexts. Several recent studies indicate that the storage and reactivation of excitatory engrams may be respectively accompanied by creation and modulation of matched inhibitory engrams. Such inhibitory engrams, also termed "negative images" or "inibitory representations," can be constructed in neural networks through simple, evolutionarily primitive, synaptic and cellular mechanisms, which are recognized to be involved in the phenomenon of excitatory-inhibitory balance observed across the brain.

Neurons and neural circuits normally operate within a preset range of activity. Outside this range, altered levels of neuronal spiking trigger a variety of homeostatic mechanisms ranging from compensatory ion-channel expression to local synaptic scaling (Nelson and Turrigiano, 2008; Turrigiano, 2012). These homeostatic mechanisms ensure that the activity parameters of a neuron operate around a set-point such that the spiking rates remain stable and a balance of depolarizing and hyperpolarizing currents is maintained. This ensures that a homeostatic equilibrium is preserved, where excitation and inhibition are both locally and globally balanced, despite plastic changes across neurons and synapses.

A potentially important candidate among these regulatory homeostatic mechanisms is potentiation of inhibitory synapses, which occurs under specific conditions and acts to prevent excessive levels of postsynaptic activity (D’Amour J and Froemke, 2015; Das et al., 2011; Fischer and Carew, 1993; Froemke et al., 2007; Vogels et al., 2011). An underappreciated property of this balancing process is that it naturally creates “inhibitory representations” or “negative images” of new, unbalanced excitatory patterns that arise within neural networks in response to experience (Barron et al., 2016a; Ramaswami, 2014). Here, we discuss how such inhibitory engrams created through compensatory inhibitory potentiation contribute to cognitive function, and ask how their disturbance could contribute to clinical features and transdiagnostic traits observed in neuropsychiatric conditions. In doing so, we integrate insights from diverse neurophysiological, behavioral, computational and brain imaging studies across multiple species, to extract and highlight fundamental principles of memory consolidation and recall (Barron et al., 2016a; Das et al., 2011; Kato et al., 2015; Letzkus et al., 2015; Ramaswami, 2014; Vogels et al., 2011).

## Direct evidence for restoration of EI balance through inhibitory synapse potentiation

Excitation and inhibition appear balanced in cortical neurons, both at a global (Moore and Nelson, 1998; vanVreeswijk and Sompolinsky, 1996; Wehr and Zador, 2003) and local level (Froemke et al., 2007; Okun and Lampl, 2008; Vogels and Abbott, 2009). At steady state, cortical neurons therefore show closely matched depolarising and hyperpolarising currents (Okun and Lampl, 2008; Wehr and Zador, 2003). However, this balance is disturbed during learning when excitatory plasticity occurs following transient reduction in inhibition (Vogels and Abbott, 2009) (Chen et al., 2015; Kruglikov and Rudy, 2008; Letzkus et al., 2015). For subsequent storage of learned information, the balance between excitation and inhibition must therefore be restored. One means by which this is achieved is via compensatory inhibitory synaptic potentation (Froemke et al., 2007; Kuchibhotla et al., 2016) (Figure 1).

**Figure 1:**
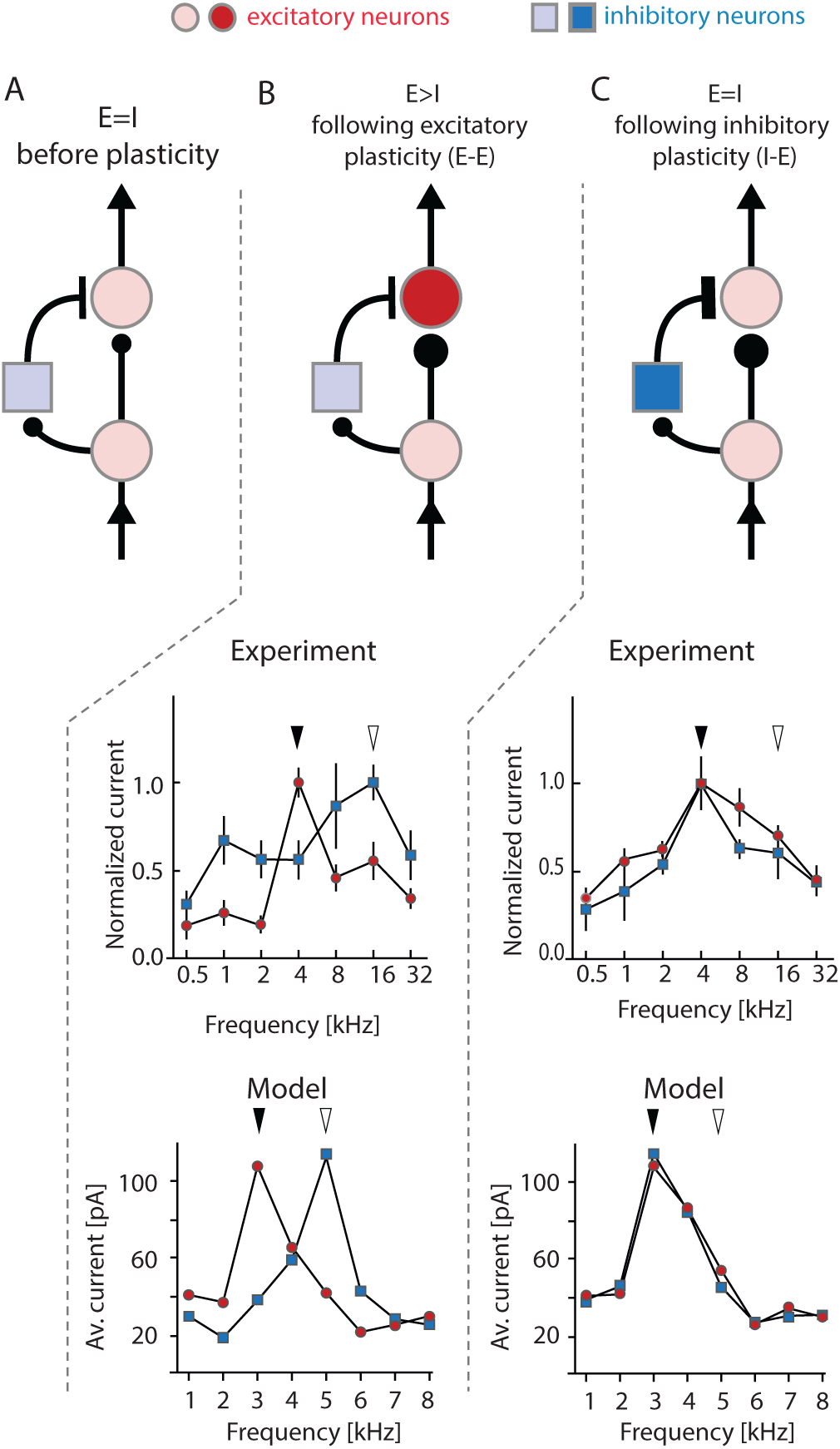
An EI rebalancing process observed in rat auditory cortex can be computationally simulated by implementing inhibitory plasticity rules. **A**. A schematic canonical circuit diagram showing excitatory neurons in red and inhibitory neurons in blue. The postsynaptic excitatory neuron receives balanced excitatory and inhibitory inputs such that the inhibitory and excitatory currents may be considered to be tuned to the same frequency. **B-C**. *Top panel:* schematic circuit diagram showing the effect of plasticity, initially leading to EI imbalance following excitatory plasticity (**B**), and subsequent restoration of balance following inhibitory plasticity (**C**). **B-C.** *Middle panel: In-vivo* whole-cell recording in primary auditory cortex with EPSCs shown in red and IPSCs shown in blue. Concurrent cholinergic stimulation and exposure to a tone (black arrow) that is shifted relative to the original preferred frequency of the neuron (white arrow), modifies the EPSC such that it retunes and peaks at the frequency of the exposed tone (**B**). After repetitive tonal stimulation (in the absence of cholinergic activity) the IPSC eventually shifts from the original to the new frequency to matches and rebalances the modified EPSC (**C**). **B-C.** *Lower panel:* Simulation from a neural network model where the excitatory tuning curve (red circles) is manually changed and a spike-time dependent inhibitory plasticity rule then applied. After 30 minutes of simulation the inhibitory tuning curve (blue squares) still shows the original tuning (**B**), however after 180 minutes it has shifted to match and rebalance the excitatory tuning (**C**). Data panels are adapted from Froemke et al., 2007andVogels et al, 2011.

At an electrophysiological level, this process of compensatory inhibitory synaptic potentiation has been best characterized in the mammalian auditory cortex. Principal cells in the auditory cortex normally show balanced depolarising and hyperpolarising currents across a range of tonal frequencies but each cell is optimally tuned to a preferred frequency at which maximal excitatory and inhibitory postsynaptic currents (EPSCs and IPSCs) occur (Wehr and Zador, 2003). Following a simple form of learning where exposure to a specific tone is paired with direct stimulation of nucleus basalis (NB), a structure that mediates attentional engagement, a shift in the preferred frequency of a principal neuron towards the frequency of the exposed tone can be observed (Froemke et al., 2007) (Figure 1b). Initial changes during this “representational plasticity” occur during NB stimulation due to the reduction in feedforward inhibition following acetlycholine release (Chen et al., 2015; Sur et al., 2013). Therefore, a shift in the preferred tonal frequency is first observed in the excitatory post-synaptic currents (EPSCs), disrupting the balance between excitation and inhibition at the new frequency (Froemke et al., 2007). Strikingly, if tone exposure is continued for 120-180 minutes without associated NB-stimulation, then a rebalancing process occurs. This rebalancing can be attributed to potentiation of the IPSC, which shifts to match the EPSC with a peak at the new frequency (Figure 1c). The mechanism responsible for this EI-balance restoration has since been investigated by both computational and experimental approaches.

Computational simulations indicate that simple synaptic plasticity rules are sufficient to account for rebalancing. In a model network of postsynaptic, integrate and fire neurons, Vogels et al. applied a simple spike-time dependent plasticity rule that acts on inhibitory synapses to mediate inhibitory-to-excitatory connections (Vogels et al., 2011). The plasticity rule allows the strength of inhibitory synapses to increase if inhibitory neurons fire within 40 msec of their excitatory postsynaptic cell, and vice versa. When this rule is implemented within an unbalanced systems where excitatory synapses are stronger than their inhibitory counterparts, simulations show that inhibitory synaptic potentiation occurs until EPSCs and IPSCs are precisely matched. With some tuning of the target spiking rate for the postsynaptic neuron, the learning rate and the spiking frequency of inhibitory neurons, the experimentally observed phenomenon of EI rebalancing can be accurately reproduced (Vogels et al., 2011) (Figure 1c).

Experimental observations and theoretical models therefore agree that inhibitory potentiation plays a critical role in rebalancing cortical networks following excitatory plasticity. At an empirical level, inhibitory potentiation is relatively poorly characterised and underlying mechanisms could vary across brain regions and inhibitory cell types. Moreover, alternative strategies, such as changes in excitatory drive (Banerjee et al., 2016), may also contribute to cortical rebalancing. Similarly, at a theoretical level, the precise learning rule responsible for inhibitory potentiation remains open to debate (Vogels et al., Frontiers 2013) as the implementation of alternative synaptic learning rules can also successfully account for cortical rebalancing (Luz and Shamir, 2012). Nevertheless, empirical investigations in rodent auditory cortex emphasise the importance of inhibitory potentiation as a means to rebalance neural cirucits, and show that near coincident pre- and post-synaptic activity is critical to the learning rule (D’Amour J and Froemke, 2015). Furthermore, inhibitory potentation is reported to be dependent upon NMDA receptors, suggesting that NMDA receptors act as coincidence detectors to trigger retrograde signals to active presynaptic neurons, thereby coordinating plasticity between co-active excitatory and inhibitory neurons (Nugent and Kauer, 2008; Nugent et al., 2007; Sudhakaran et al., 2012; Woodin et al., 2003). Thus, by simulating biologically plausible learning rules for inhibitory potentiation, theoretical models can explain key features of EI balancing observed during representational plasticity in the auditory cortex (Figure 1c).

## Negative images in adaptive stimulus filtering and behavioral habituation

Following increased activity across an ensemble of excitatory synapses, the form of EI balancing described above is predicted to result in delayed strengthening of matched inhibitory synapse ensembles (Barron et al., 2016a; Ramaswami, 2014) (Figure 2A-C). This creates inhibitory engrams or representations, which counterbalance excitatory representations and ensure that EI balance is maintained in face of increased excitation. In large-scale neural networks, such inhibitory engrams may underlie multiple fundamental cognitive processes. We first consider their role in habituation, a form of non-associative, implicit memory that reduces innate responses to irrelevant stimuli (Ramaswami, 2014; Rankin et al., 2009; Thompson and Spencer, 1966).

**Figure 2:**
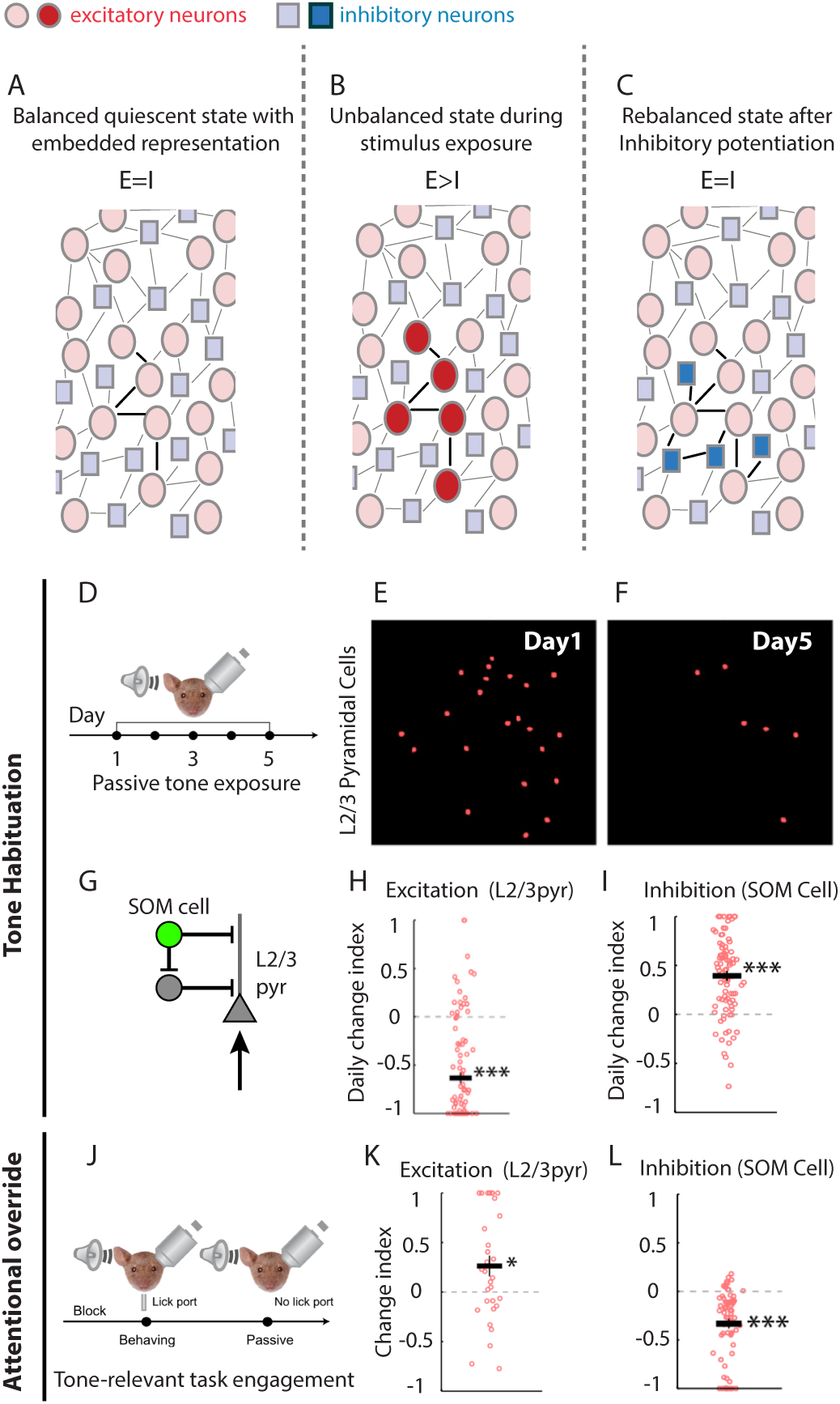
Inhibitory representations can mediate sensory habituation. **A-C.** Schematic model for habituation based on (Ramaswami, 2014). **A**. A balanced circuit of inhibitory (blue) neurons and excitatory (red) neurons showing a connected excitatory representation. **B**. Continuous exposure to the relevant percept for the excitatory representation referred to in *A* results in EI imbalance, due to sustained excitatory activity (bright red). **C**. EI rebalancing through inhibitory potentiation creates a matched inhibitory representation (bright blue) to suppress the excitatory representation. **D-I.** Direct evidence for inhibitory potentiation during habituation from mouse auditory cortex (adapted from Kato et al. 2015). **D**. Habituation protocol: mice were passively exposed to a tone for 5 days. Mice experienced 200 trials per day and on each trial the tone lasted for 5 to 9 seconds. **E-F**. In-vivo two-photon image of tone-evoked GCaMP6s-expressing neurons in layer 2/3 of auditory cortex on day 1 (**E**), and on day 5 (**F**). Excitation was sparser day 5. **G.** Canonical cortical microcircuit showing SOM-positive inhibitory neurons connecting to layer 2/3 (L2/3) excitatory neurons. **H-I.** The daily change index, used to assess the change in activity across days 1-5, showed a reduced in excitation in layers 2/3 (**H**) which was accompanied by an increase in excitation in SOM-positive inhibitory neurons (**I**). Together these results suggest a reduction in excitation and an increase in inhibition following habituation. **J-L.** In habituated animals, tone-relevant task engagement is accompanied by disinhibition of L2/3 excitatory cells (adapted from Kato et al. 2015). **J**. Tone-relevant task protocol: Habituated animals were trained to lick a food port in response to the habituated tone. The response was then compared to passive exposure to the habituated tone, in the absence of the food port. **K-L.** The change index reflects the increase in the layer 2/3 neuron excitation during the tone-relevant task, relative to the passive exposure. A task associated increase in the excitatory response was observed (**K**), which was accompanied by a decrease in tone-evoked activity in SOM-positive neurons (**L**), consistent with attentional disinhibition.

To explain olfactory habituation in *Drosophila,* behavioral genetic analyses have inferred and invoked an inhibitory learning rule similar to that involved in restoring EI balance in mammalian auditory cortex, (Das et al., 2011; Larkin et al., 2010; Ramaswami, 2014). In insects, odorants are encoded by assemblies of projection neurons in the antennal lobe, a structure homologous to the mammalian olfactory bulb. Many lines of evidence argue that in a neutral environment, prolonged activation of an odorant-specific excitatory assembly results in the selective strengthening of inhibitory synapses onto neurons activated by the odorant (Das et al., 2011; Glanzman, 2011). This results in the formation of an inhibitory replica of the specific pattern of odor-induced excitation. The newly created inhibitory representation of odorant-induced excitation acts as a filter to specifically attenuate physiological and behavioural responses to the familar and inconsequential odorant (Ramaswami, 2014). Significantly, as observed for the EI balancing process associated with re-tuning cells in mammalian auditory cortex, insect olfactory habituation also requires postsynaptic NMDA receptors suggesting that a common homeostatic mechanism is at play (D’Amour J and Froemke, 2015; Das et al., 2011). Together, these observations suggest that olfactory habituation may be usefully considered to arise through a form of EI balancing in which inhibitory potentiation serves to restrain the spiking of a subset of odorant-activated projection neurons to within a behaviorally appropriate range.

Broadly similar mechanisms for auditory habituation have been recently reported in the mammalian cortex, albeit without direct evidence for the underlying synaptic mechanism (Kato et al., 2015) (Fig 2D-L). Here, *in vivo* GCaMP-based imaging shows that tone-specific auditory habituation is associated with reduced calcium-fluxes in layer 2/3 pyramidal cells in the rodent auditory cortex. This reduction in pyramidal cell activity is accompanied by enhanced activity of somatostatin positive (SOM) inhibitory neurons in the same brain region. Habituation can thus be characterised as a 10-fold reduction in the excitation/inhibition ratio of population activity. Interestingly, when attention to the tone becomes important for task performance, SOM inhibition is reduced and pyramidal cell responses to the tone increase even in habituated animals (Kato et al., 2015).

Taken together, data from insect and mammalian nervous systems suggest that habituation may generally arise through the formation of inhibitory engrams created by inhibitory synaptic potentiation via mechanisms that resemble those involved in EI balancing (D’Amour J and Froemke, 2015; Froemke et al., 2007; Kato et al., 2015; Ramaswami, 2014; Vogels et al., 2011). We note that in the mammalian brain, the subtypes of inhibitory interneurons involved and their mode of regulation may depend on the particular neural circuit in which they are embedded. For instance, while SOM-positive interneurons have been identified as playing a critical role in auditory habituation, PV-positive interneurons are noted for their more pervasive role in equalising the ratio between inhibition and excitation (Xue et al., 2014). Although the precise interaction between these different subtypes is not yet clear, the model in which matched inhibition drives habituation generates important predictions. One notable corollary suggests that innate behavioural responses attenuated by habituation can later be rapidly restored through disinhibition, via inputs that silence the relevant inhibitory neurons (Kato et al., 2015). This provides an explanation for the psychological phenomena of dishabituation and attentional override, two defining properties of habituation (Ramaswami, 2014; Rankin et al., 2009; Thompson and Spencer, 1966).

## Regulating the formation of inhibitory engrams

As evidenced from studies of representational plasticity and habituation discussed above, both the formation and activity of inhibitory engrams are regulated by behavioral context. But under what circumstances do these inhibitory engrams form? By definition, behavioural habituation occurs to non-salient and inconsequential stimuli. Indeed, contextual inputs that confer salience or assign emotional significance to a stimulus are known to actively block habituation (Ramaswami, 2014; Rankin et al., 2009; Thompson and Spencer, 1966). This is probably best revealed by physiological studies investigating regulation of intrinsic inhibitory synaptic plasticity (metaplasticity) in the circuit that underlies habituation of a siphon withdrawal reflex in Aplysia (Bristol and Carew, 2005; Fischer et al., 1997). When multiple mild stimuli are applied to the tail, a reduced tail-touch induced siphon withdrawal is observed as a consequence of potentiated inhibitory feedback onto siphon motorneurons. By contrast, a single electric shock applied to the tail increases tail-touch induced sensitized siphon withdrawal through serotonin-dependent potentiation of excitatory sensorimotor synapses. Remarkably, the tail-shock induced release of serotonin not only potentiates excitatory connections but also blocks habituation by reducing the ability of repeated, mild tactile tail stimulation to cause inhibitory interneuron-motorneuron synapse facilitation (Bristol and Carew, 2005; Fischer et al., 1997) (Figure 3).

**Figure 3:**
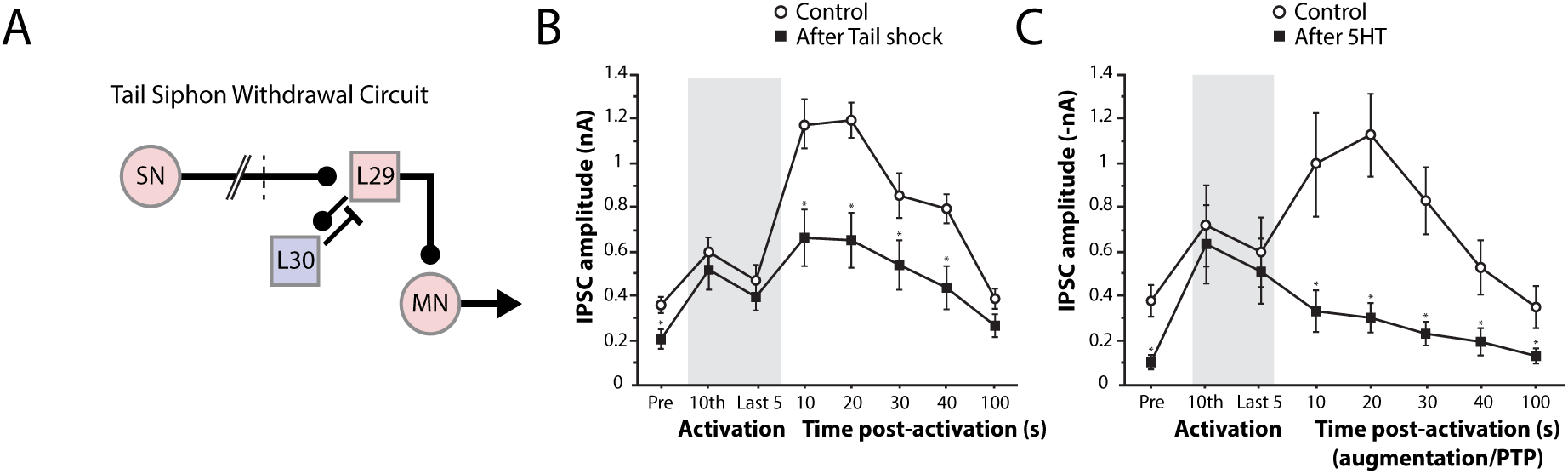
Neuromodulation gates inhibitory potentiation and EI balancing. (Adapted fromFischer et al., 1997). **A**. Simplified circuitry that mediates the tail-siphon withdrawal (T-SW) response in *Aplysia.* Sensory neurons (SN) in the tail activate an excitatory interneuron L29 (shown in pink) which excites both the siphon motor neuron (MN) and the inhibitory interneuron L30 (shown in blue). L30 mediates feedback inhibition onto L29. Tail shock results in serotonin release on both L30-L29 and sensorimotor synapses. T-SW habituation is accompanied by enhanced L30-L29 transmission (increased IPSCs recorded in L29), which can also be induced by direct L30 stimulation at frequencies normally induced by tail touch (Fischer, 1997; Bristol et al., 2005). **B-C**. Brief tetanic stimulation was delivered to the inhibitory interneuron L30 (shaded bar indicates stimulation period). The IPSC was recorded from the excitatory interneuron L29. Following stimulation, the IPSC from L29 was enhanced (control condition, open circles). However, this enhancement was not observed if the tetanic stimulation to the inhibitory interneuron L30 was delivered 90 seconds after a tail-shock (**B**, filled squares, * indicates a significant difference of p<0.05 relative to the control condition). Furthermore, this enhancement was also not observed if the tetanic stimulation to the inhibitory interneuron L30 was delivered 90 seconds after application of serotonin (**C**, filled squares, * indicates a significant difference of p<0.05 relative to the control condition). Together these results suggest that inhibitory potentiation is similarly blocked by either tail-shock or direct application of serotonin. In other scenarios, neuromodulators could act by prevent inhibitory neuron firing.

Interestingly, in the mammalian auditory cortex, inhibitory potentiation following EPSC enhancement occurs in response to continued tonal stimulation, but this has only been observed in the absence of cholinergic NB stimulation (Froemke et al., 2007). As cholinergic NB afferents mediate disinhibition (Froemke, 2007), this leads to the prediction that concurrent NB stimulation would actively inhibit heterosynaptic inhibitory plasticity to prevent EI rebalancing. More generally, we speculate that when memories are actively encoded during transient periods of high neuromodulator concentration, EI rebalancing mechanisms may be disrupted. Abnormal persistence of such neuromodulation may result in so-called maladaptive memories that persistently reactivate, as observed in post-traumatic stress disorder (Ehlers and Clark, 2000). In conclusion, processes that underlie the restoration of EI balance in both the retuning of cells in the mammalian auditory cortex and in behavioural habituation best described in invertebrates, not only both rely on inhibitory plasticity, but may also be regulated in a similar manner by context-dependent neuromodulation.

## Inhibitory engrams in associative memory storage and recall

In addition to the proposed role for inhibitory potentiation in cellular re-tuning and habituation, recent experiments in humans suggest that inhibitory potentiation may also play a fundamental role in the storage and recall of associative memories (Barron et al., 2016a) (Figure 4). Using representational fMRI (Barron et al., 2016b), the expression of associative memories can be indexed immediately after learning, within the region of cortex that typically represents the stimuli in question. For example, when participants learn to associate rotationally-invariant abstract shapes, representational overlap is observed in an anterior region of the lateral occipital cortex (LOC). However, over time, this cortical expression of the associative memories decreases, such that it becomes invisible to fMRI 24 hours after learning. Remarkably, when the level of the cortical inhibitory transmitter GABA is reduced using anodal transcranial direct current stimulation (tDCS), the expression of the associative memory reappears. Thus, associative memory transitions from an early form which is visible to fMRI, to a later quiescent state that can only be revealed under conditions of reduced inhibition. This result is consistent with the idea that excitatory synaptic potentiation that occurs during learning, is later matched by equivalent inhibitory synaptic potentiation and the formation of a matching inhibitory representation.

**Figure 4:**
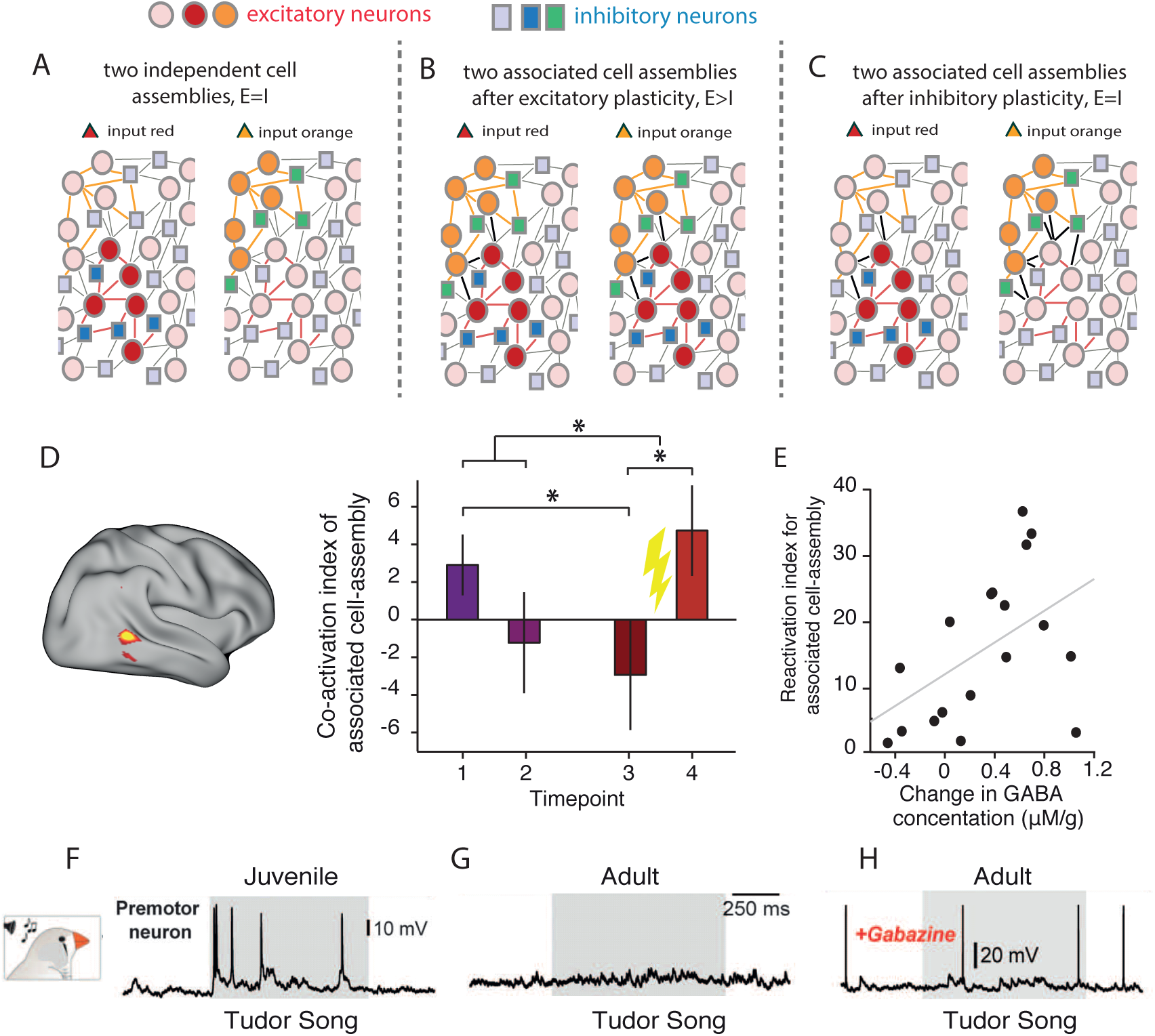
Inhibitory rebalancing allows associative information to lie dormant unless EI balance is disturbed. **A-C.** Schematic representation of a neural network that stores two distinct cell assemblies which are represented by an ensemble of excitatory (either red or orange circles) and inhibitory (either blue or green squares). Initially the two cell assemblies are distinct, with one responding to the red input, and the other responding to the orange input (**A**). When the two cell assemblies become associated via excitatory plasticity between the two excitatory ensembles (black lines), co-activation is observed such that the red cell assembly responds to the orange input and the orange cell assembly responds to the red input (**B**). If inhibitory plasticity then acts to restore balance in the network, the newly strengthened excitatory connections are quenched by inhibition such that co-activation between the two cell assemblies is reduced (**C**). **D.** In the human brain, representational fMRI can be used to index co-activation of neural representations for associated abstract shapes in an anterior region of the lateral occipital complex. However, this co-activation is only observed immediately after learning (timepoint 1), and decreases over time (timepoint 2 (~1 hour after learning) and timepoint 3 (~24 hours after learning)). Once the associative memory is quiescent, application of anodal transcranial direct current stimulation leads to a significant reduction in cortical GABA and an increase in the co-activation index for associated stimuli. This suggests that associative memories are stored in cortex in balanced excitatory-inhibitory ensembles, which lie dormant unless the balance between excitation and inhibition is disrupted. **E.** The increase in the co-activation index between associated representations can be predicted by the drop in GABA induced by brain stimulation. **F.** In juvenile songbirds, premotor neurons show an increase in activity during exposure to a tutor song. **G.** In adult songbirds, premotor neurons are quiescent during exposure to a tutor song. **H.** Application of a GABA agonist, Gabazine, releases excitation in premotor neurons of the songbird, revealing a response not dissimilar from that observed in developing juvenile birds.

Together these observations point to the following model (Figure 4). Excitatory potentiation during learning enhances connectivity between distinct neuronal ensembles that encode the associated stimuli in question. EI balance is then restored by inhibitory synaptic potentiation, which at a network level creates an inhibitory engram of the newly strengthened excitatory connections to match and cancel/ reduce their effect. Once the balanced state is restored, weakly connected excitatory neurons are balanced by weak inhibition, while strongly connected excitatory neurons are balanced by strong inhibition. Thus, strong associations are stored in a latent state that allows them to be selectively revealed by disinhibition (Barron et al., 2016a).

These findings are consistent with electrophysiological recording in a songbird while it listens to a tutor’s song (Figure 4). When juvenile zebra finches are exposed to a tutor song, excitatory neurons in HVC are active. However, in adult birds HVC-neuron responses to tutor song are silenced by inhibitory inputs, whose strength is proportional to the relative similarlity of the birds own song to the tutors (Vallentin et al., 2016). Disinhibiting the HVC via application of a GABA agonist, Gabazine, releases excitation to reveal a response not dissimilar from that observed in developing juvenile birds. This suggests that inhibition plays an important role in protecting stored representations from interference with closely related information.

The temporal period and the precise mechanisms involved in the inferred EI balancing process are not known. But a particularly attractive notion is that this occurs during sleep, potentially during sharp-wave ripples, when previous experiences and memories are replayed offline (Eschenko et al., 2008; Ji and Wilson, 2007; Ramadan et al., 2009). We speculate that this could be one of the ways in which sleep contributes not only to memory consolidation, but also to homeostatic processes that prepare the brain for new learning.

Within this framework, it remains unproven as to how stored information is normally recalled. While mechanisms for memory recall could vary across brain systems and circuits, we suggest cortical disinhibition as one potential strategy to allow release of strongly connected excitatory ensembles from balanced strong inhibition (Kato et al., 2015; Vallentin, 2016; Barron et al., 2016a). Disinhibition may be mediated by neuromodulators, acting for instance through muscarinic acetylcholine receptors on cortical interneurons (Kim et al., 2016). More selective disinhibition may be additionally mediated through neural structures critical for memory recall, such as the hippocampus. Thus, in addition to the established function of local disinhibition to enhance excitatory transmission (Vogels and Abbott, 2009) and contribute to the initial encoding of memory (Chen et al., 2015; Kruglikov and Rudy, 2008; Letzkus et al., 2015), disinhibition may also play a second significant function to facilitate release of previously learned, but latent cortical associations (Barron et al., 2016a).

Experimental evidence for disinhibition in memory recall is particularly evident following fear conditioning, the most simple form of associative memory (Courtin et al., 2014; Letzkus et al., 2015; Wolff et al., 2014). Here, when a conditioned auditory stimulus (CS) is used to trigger fear-memory recall in rats, phasic inhibition of a subset of parvalbumin-positive (PV) GABAergic interneurons can be observed in dorso-medial prefrontal cortex (Courtin et al., 2014). This higher level inhibition of inhibitory neurons can be conceived to bring about disinhibition of the learned fear response. Indeed, optogenetic inhibition of these GABAergic neurons is sufficient to induce freezing (Courtin et al., 2014). This disinhibition may be mediated by vasointestinal peptide positive (VIP) interneurons (David et al., 2007; Pfeffer et al., 2013; Pi et al., 2013), although details of the disinhibitory elements and their circuitry can vary between different brain regions and types of associative learning (Fu et al., 2014; Wolff et al., 2014).

## A framework for normal and variant memory storage and recall

The observations and arguments above lead to a common systems level framework for memory storage and recall that are built on two strikingly simple principles (Figure 5). (1) Following habituation or associative training respectively, innate behaviors and simple associative memories are masked by inhibitory engrams. These compensatory inhibitory representations ensure that EI balance is maintained despite new learning and may be considered a critical component of memory consolidation. During habituation, inhibitory engrams are created when stimuli are experienced without concurrent engagement of emotional or attentional circuits (Figures 1-2, Figure 5)._Following associative learning, we speculate that sharp-wave ripple dependent replay during sleep may play an important role in their formation. (2) Innate behaviours and associative memories can be recalled in appropriate contexts through selective and local disinhibition. We speculate that local disinhibition is driven by contextual or attentional inputs that recruit secondary inhibitory neurons to selectively target cell-types encoding inhibitory engrams (Chen et al., 2015; Letzkus et al., 2015; Perisse et al., 2016) (Figure 2; Figure 4; Figure 5). In this manner volitional and directed recall of memories may occur via sensory stimulation or attentional mechanisms, which act to disinhibit and activate cortical subdomains that store specific memories. Although we focus here on inhibitory engrams created by local EI balancing mechanisms, the framework we propose also naturally accommodates inhibitory representations constructed through potential alternative mechanisms (Clark, 2013; Cooke et al., 2015; Kogo and Trengove, 2015; Ramaswami, 2014; Rao and Ballard, 1999; Wacongne et al., 2012).

**Figure 5:**
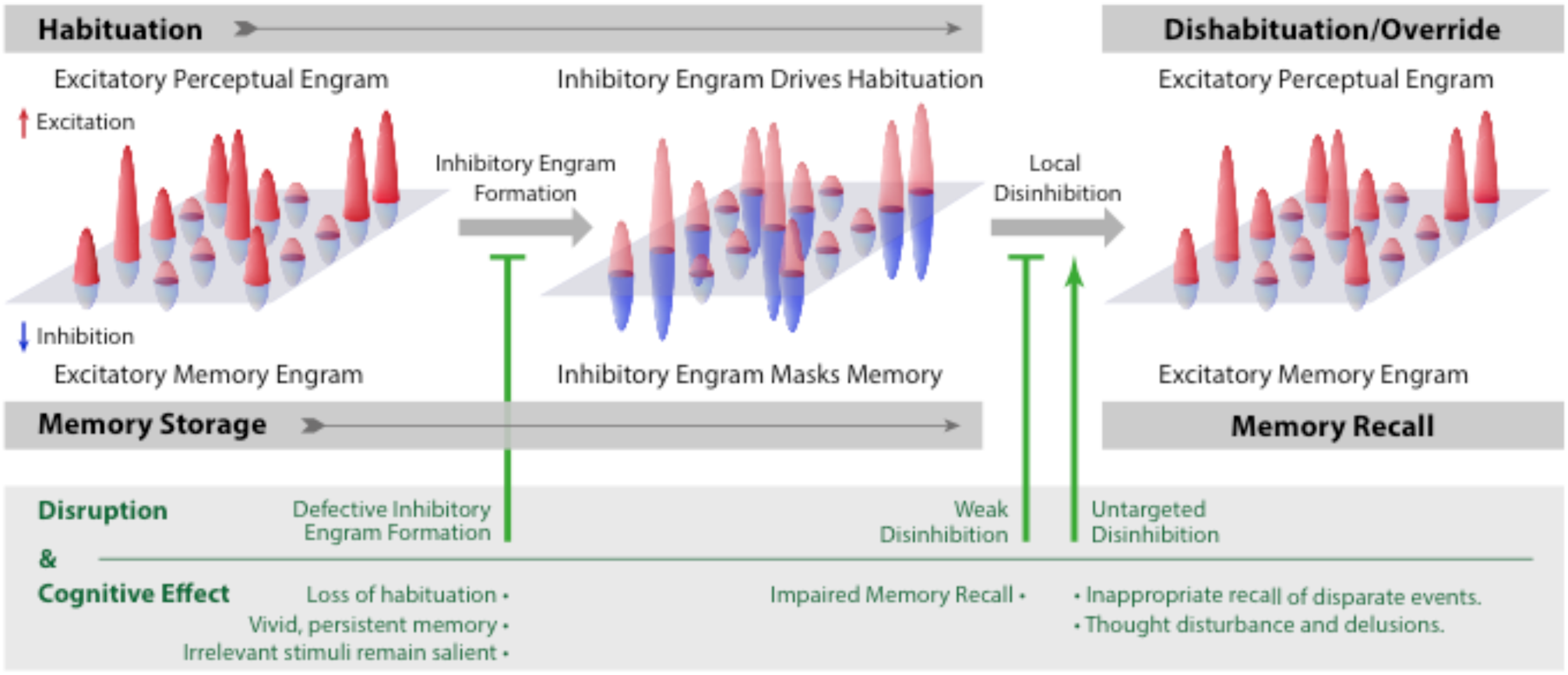
Inhibitory engrams in perception and memory: a model. A hypothetical framework showing how inhibitory engrams could form in neural systems illustrated with reference to their influence in perceptual habituation, memory storage and recall. Excitatory inputs onto an array of postsynaptic neurons are shown as positive red peaks and inhibitory inputs as negative blue peaks. In the illustration above, these constitute excitatory and inhibitory engrams respectively. Before habituation, sensory stimuli activate excitatory perceptual engrams but trigger relatively weak or imprecise inhibition. Repeated excitatory engram stimulation, with minimal attentional or emotional engagement, results in formation of a matched inhibitory engram that reduces the stimulus response and causes behavioral habituation (Ramaswami, 2014). Attentional inputs or dishabituating stimuli can restore the initial stimulus response by promoting disinhibitory inputs that suppress inhibitory engrams (Ramaswami, 2014; Kato et al., 2015). Similarly, memory is first encoded as excitatory engrams. Over time, matched inhibitory engrams form to rebalance the neural network (Barron et al., 2016; Valentin et al., 2016). The formation of these inhibitory memory engrams may occur via homeostatic mechanisms similar to those involved in habituation, potentially during replay of excitatory memory engrams. Once formed, inhibitory engrams allow memories to be stored in a dormant form for context-appropriate recall, which we hypothesise to occur through focussed disinhibition. (Below in green) Defects in inhibitory engram formation or regulation are predicted to cause perceptual and cognitive abnormalities including transdiagnostic traits observed in autism, schizophrenia and post-traumatic stress disorders.

The model appears to have considerable explanatory and heuristic value. For instance, it conceptually distinguishes between behavioral amnesia arising from defects in storage from those arising from defects in recall. It is therefore relevant to recent work which shows that a memory engram remains preserved in amnestic rats that do not express contextual fear memory (Roy et al., 2016; Ryan et al., 2015). Consistent with the framework we present here, in the mouse models of Alzheimer’s disease, memory storage largely remains intact while recall is disrupted, potentially by a failure of disinhibitory pathways that mediate recall of stored memories (Figure 5).

## Implications for EI disruption in clinical conditions

Many investigators have worked towards establishing a common framework to account for the behavioural phenotypes common to psychiatric conditions, which otherwise have complex and distinct genetic bases (Lisman et al., 2008; Markram et al., 2007; Oberman et al., 2005; Sahin and Sur, 2015; Walsh et al., 2008; Yizhar et al., 2011). Our premise that a major function of EI balancing is to mask irrelevant perceptions, memories and behavior, allows mechanistic predictions to be drawn that concern specific cognitive dysfunctions that may arise when the process is disrupted (Figure 5). These predictions are particularly relevant, given several lines of genetic and physiological evidence pointing to an imbalance between excitation and inhibition as a possible substrate for symptoms observed in a range of clinical conditions including autism and schizophrenia (Chou et al., 2010; Gogolla et al., 2009; Han et al., 2014; Lisman et al., 2008; Nelson and Valakh, 2015; Penagarikano et al., 2011; Rubenstein and Merzenich, 2003; Tuchman et al., 2010; Tyzio et al., 2014; Yizhar et al., 2011).

One predicted consequence of defects in the formation of inhibitory engrams is weak behavioral habituation that could arise either from increased neuromodulatory activity which blocks the formation of inhibitory representations, or from defects in inhibitory function and plasticity. Remarkably, weak habituation is particularly common feature of autism spectrum disorders (ASD) that is also observed in schizophrenia (Ethridge et al., 2016; Guiraud et al., 2011; Kleinhans et al., 2009; Rankin et al., 2009; Sinha et al., 2014). Weak habituation could also result in several downstream cognitive changes observed in autism and in schizophrenia, including altered sensory gating, stimulus hypersensitivity and reduced ability to cope in complex environments, where multiple signals may appear salient and compete for attention (Ethridge et al., 2016; Ramaswami, 2014; Rankin et al., 2009; Sinha et al., 2014; Vogels and Abbott, 2007). It is also conceivable that additional features of autism such as sticky attention, wherein familar stimuli remain engaging for unusually long periods of time while novel stimuli seem challenging, could arise from weak habituation and hypersensitivity to novel stimuli (Landry and Bryson, 2004; Sinha et al., 2014; Vogels and Abbott, 2007). An additional predicted consequence of defects in inhibitory rebalancing that we postulate to be necessary for masking associative memories, is strong and contextually inappropriate recall of associative memories. Strikingly, unusually strong and vivid associative memories are often associated with ASD (Zamoscik et al., 2016). It is conceivable that vivid recall of unconnected memory traces could lead to inappropriate associations and thought disturbances and delusions observed in schizophrenia (Figure 5). This is consistent with theoretical models in which EI imbalance in neural networks has been used to account for hallucinations and delusional experiences that may be observed both in autism and in schizophrenia (Gao and Penzes, 2015; Toal et al., 2009; Vogels and Abbott, 2007).

This mechanistic proposal that specific cognitive features and transdiagnostic traits of autism and schizophrenia arise in part due to defects in the creation of inhibitory engrams is now experimentally testable, taking advantage of MRI paradigms that provide an index for EI balance across associative memories (Barron et al., 2016b; Barron et al., 2016a). One simple testable prediction is that EI imbalance across cortical associations will persist for unusally long time periods in affected individuals. While the validity of this proposal will require additional investigation, we suggest that a model that connects synaptic mechanisms of EI balancing in neural circuits with fundamental cognitive processes altered in psychiatric conditions may provide a unifying principle to explain common disturbances that arise from diverse molecular and genetic perturbations.

## Conclusion

In summary, the model that inhibitory engrams created by EI balancing mechanisms can reversibly suppress and thereby allow context-relevant recall of innate and learned behaviors provides an elegant mechanistic framework to understand fundamental cognitive functions in both health and disease. Specifically, this framework provides a simple description for key processes that underlie memory storage and recall in psychotypical cognitive function, which gives insight into the substrates of neuropsychiatric symptoms that emerge when EI balance is disturbed.

## AKNOWLEDGEMENTS

We thank Jens Hillebrand for help with the figures, Isabell Twick for comments on the manuscript, Kevin Mitchell, Srikanth Ramaswamy and Mriganka Sur for useful discussions. H.C.B. is supported by a Junior Research Fellowship from Merton College, University of Oxford, and the John Fell Oxford University Press (OUP) Research Fund (153/046). T.P.V. is funded by a Sir Henry Dale Fellowship from the Wellcome Trust (WT100000). T.E.J.B. is supported by a Wellcome Trust Senior Research Fellowship (WT104765MA) together with a James S. McDonnell Foundation Award (JSMF220020372). M.R. acknowledges grants from Science Foundation Ireland and the Simons Foundation Autism Research Initiative as well as core support from the NCBS, Tata Institute of Fundamental Research, Bangalore for collaborative work with K. VijayRaghavan.

